# Efficient and durable gene activation by Cas9-mediated epigenome editing in vivo

**DOI:** 10.1101/2024.05.03.592438

**Authors:** Yuanbo Qin, Talha Akbulut, Rajakumar Mandraju, Keith Connolly, John Bechill, Farzaneh Assadian, Alison Shottek, Seth Levy, Jamie Benoit, Tetsuya Yamagata

**Affiliations:** Modalis Therapeutics, Inc., Waltham, Massachusetts, USA

## Abstract

Epigenome editing technology holds great promise for treating diverse genetic disorders. While a series of advances has been made on epigenetic silencing using programmable editors, little progress has been made in leveraging epigenetic activation for therapeutic application. Here we demonstrate epigenetic activation of the LAMA1 gene for the treatment of LAMA2-CMD, a severe congenital muscle dystrophy (CMD) caused by biallelic mutations in the LAMA2 gene. LAMA1 is a sister homologue that is known to compensate for the function of LAMA2. However, supplementing LAMA1 or LAMA2 gene via viral platform is not feasible due to the large size of their coding sequences. Through a single administration of our (Adeno-associated virus) AAV vector encoding all the necessary elements for epigenetic activation, a platform termed CRISPR guide-nucleotide directed modulation (GNDM), we observed significant LAMA1 gene upregulation and phenotype improvements in DyW mice, a severe disease model of LAMA2-CMD. Notably, sustained expression of the GNDM gene and subsequent activation of the LAMA1 gene persisted beyond analyzed period of one year despite immune recognition of the GNDM protein by the host immune system. Regulatory T (Treg) cells appeared to facilitate tolerance to GNDM in the transduced muscle tissue. The muscle-tropic AAV capsid exhibited desired vector biodistribution and promising pharmacodynamics with good safety profiles in adult non-human primates (NHPs). Moreover, administration to juvenile NHPs demonstrated superior pharmacodynamics compared to adults, even at half the adult dose, suggesting safer and more effective therapeutic outcomes in mostly pediatric LAMA2-CMD patients. Our approach holds broad applicability for a range of loss-of-function genetic disorders and could offer a therapeutic breakthrough where active epigenome brings clinical benefit.

## Introduction

Epigenome editing has emerged as a promising new strategy for activating or silencing genes without altering their DNA sequence^1^. In this context, the epigenetic modifier domain is fused with a programmable DNA-binding domain (DBD) to direct its effect towards the designated regulatory element of a target gene^2, 3^. For epigenetic activation, a selected DBD is fused with an engineered activation moiety to activate target genes. While a series of efforts has been made for epigenetic silencing towards various therapeutics application^4–11^, little progress has been made on epigenetic gene activation for clinical use since its conception^12–14^. This gap comes from a series of challenges associated with epigenetic activation. First, epigenetic activation needs creation of active local chromatin structures, which requires formation of an active transcriptional activation complex around the regulatory element of a target gene^15^. For this purpose, a robust DBD platform, delivered by a suitable vector, is necessary to maintain the local active chromatin state^2^. Second, gene activation involves gene- and cell-type specific machinery, and thus selection of a context-dependent activation moiety is needed for optimum epigenetic activation^16^. Third, potent gene activation moieties tend to be larger in size compared to compact silencer domains (such as KRAB), However, for practical therapeutic use all the necessary elements for intended epigenetic activation need to be within the cargo limit of a single viral vector^16, 17^. Finally, a drug candidate faces uncharted pharmacodynamic and safety profile challenges in preclinical testing because of the lack of precedence in epigenetic activation drugs. Here we tackled these issues and demonstrated, for the first time, that epigenetic activation is a safe and viable strategy for clinical application to treat a genetic disease with unmet medical needs.

LAMA2-CMD is one of the most common forms of congenital muscular dystrophy, typically presenting symptoms at birth or as early as the embryonic stage. It is characterized by severe hypotonia, progressive muscle wasting, significant impairment in daily activities, and typically, death occurs in the second decade of life^18, 19^. Currently, there is no cure or approved therapies for this disorder^20^. LAMA2-CMD is an autosomal recessive disorder caused by mutations in both copies of the LAMA2 gene, causing a deficiency of its coding protein Laminin α2. Laminin α2, together with its family subunits β1 and γ1, form a trimeric complex known as laminin-211 (merosin)^21^. Laminin-211 is a crucial component of the extracellular matrix in muscle tissue, playing a vital role in muscle fiber stability and signal transduction^20^. The lack of Laminin α2, resulting in deficiency of laminin-211, disrupts both the integrity of muscle tissue and normal muscle cell function, leading to skeletal muscle degeneration. LAMA1 codes for a functionally and structurally similar protein to LAMA2, which is expressed during embryonic development but absent in muscle tissues post-birth. Several studies have shown that LAMA1 can compensate for the loss of LAMA2 gene in animal models^22–25^. Thus, supplementing the LAMA1 or LAMA2 gene via gene therapy would be a technical solution to LAMA2-CMD.

AAV is widely used for gene therapy applications. Direct supplementation of LAMA1 or LAMA2 gene using AAV is unfeasible due to their over-sized coding sequence (> 9 kb), which exceeds the packaging capacity of AAVs. In an attempt to circumvent this problem, CRISPR-based epigenetic activation (CRISPRa) has been previously shown to successfully upregulate LAMA1 gene in a disease model of LAMA2-CMD and ameliorate the disease phenotypes^26^. However, there are some pharmaceutical hurdles mentioned earlier in advancing this concept to a viable therapeutic. To overcome the challenges, we developed CRISPR-GNDM^®^ technology, which features a compact Cas9 variant fused with engineered transcriptional modulators (collectively called GNDM) that can be packaged into a single AAV vector with high specificity and minimum off-target effects.

Here, we demonstrate GNDM-mediated upregulation of Lama1 in wildtype mice and LAMA2-CMD disease models at multiple ages and doses. Immunogenicity and pharmacodynamics were carefully examined to demonstrate the safety and efficacy of our vector. A surrogate vector was then utilized in two NHP studies to assess safety, biodistribution, immunogenicity, and pharmacodynamics in two different ages (adult and juvenile) to provide proof of concept and identify the optimal dose for our intended patient age group. To our knowledge this is the first demonstration of epigenetic activation in vivo that is durable and efficacious. This study represents one of the first demonstrations of successful systemic epigenetic editing in large animals. Additionally, our study is among the first to prove that targeted Cas9 expression by systemic delivery is safe and well-tolerated in a large animal model. These findings underscore the promise of CRISPR-GNDM^®^ technology as a next-generation gene therapy platform for a variety of neuromuscular and other genetic disorders.

## Results

### AAV9-GNDM-m31 induces Lama1 gene expression with high specificity

Therapeutic activation of LAMA1 requires continuous presence of an epigenetic activator at the correct region of the regulatory element of the LAMA1 gene. To this end, an AAV delivery system and a Cas9 DBD platform were chosen for their long-term transgene expression and robust DNA-binding capacity, respectively. Due to space limitations, a single guide (gRNA) was used in the construct. Thus, to discover the most potent gRNAs for upregulating mouse Lama1 gene expression, we transfected C2C12 mouse myoblast cells with plasmids expressing both GNDM and an arrayed library comprising 48 gRNAs spanning a 2kb region surrounding the Lama1 gene transcription start site (Figure 1A). This region was selected for the enrichment of epigenetic markers highly suggestive for a key regulatory element^27^. qPCR analysis revealed that several gRNAs significantly increased Lama1 expression compared to non-targeting gRNAs (NTGs), leading to the selection of gRNA m31 for further validation (Figure 1B).

**Figure 1:**
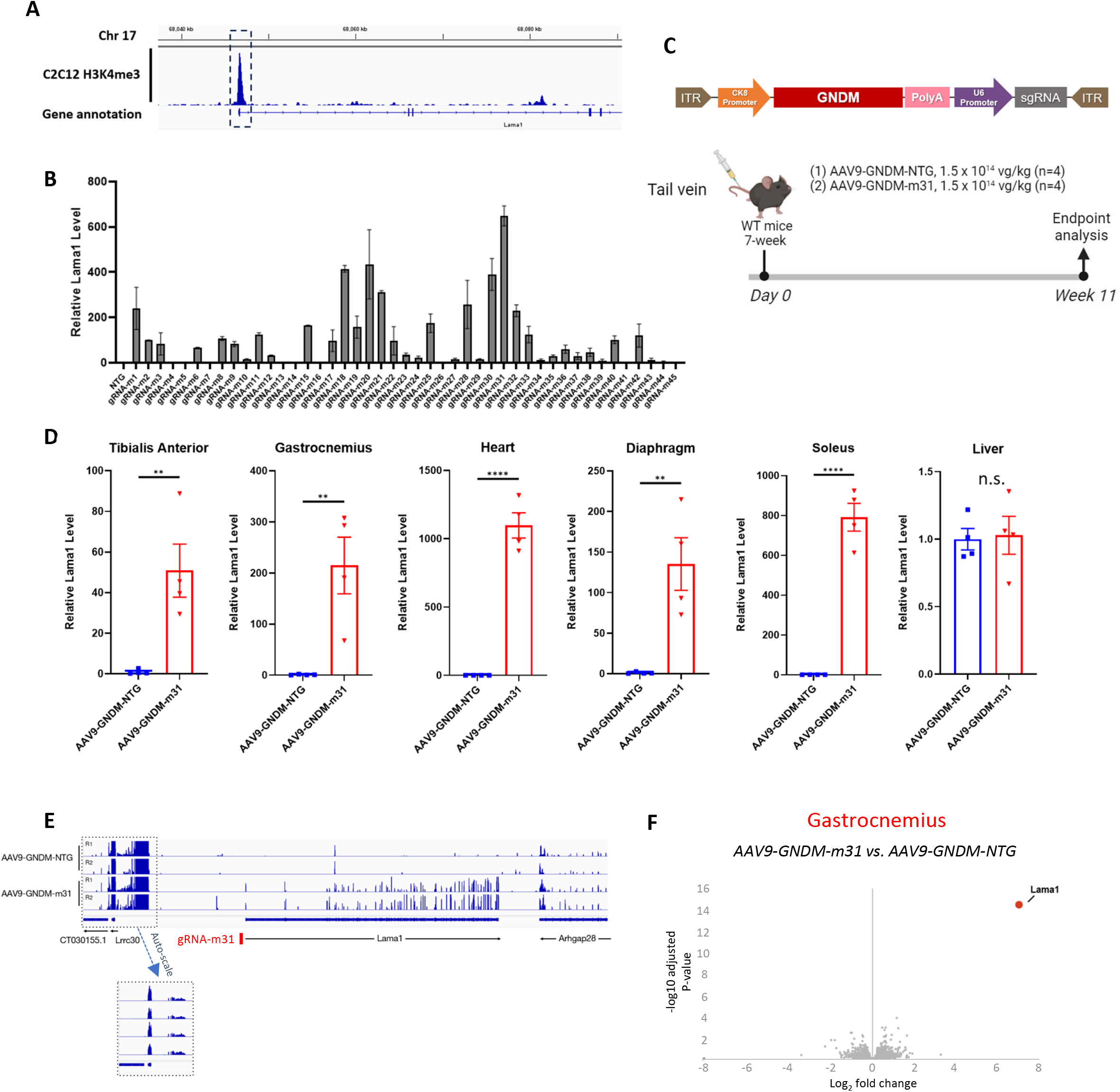
Induction of Lama1 expression via GNDM with high specificity *in vitro* and *in vivo*. (A) Mouse genomic (mm9 genome assembly) features of GNDM target site at the Lama1 locus. Tracks indicate mouse Lama1 gene annotation, chromosome 17 coordinates as well as enrichment of H3K4me histone mark in C2C12 cells determined by ChIP-seq assay. Dot-line highlighted region was selected for gRNA screening for Lama1 upregulation in C2C12 cells. (B) Summary of gRNA screening results in C2C12 cells. Relative Lama1 mRNA level is determined by QPCR where each sample expression values were normalized to the non-targeting gRNA (NTG) sample values. Bars represent mean values and SEM is represented by error bars (n = 3). (C) A schematic of the GNDM transgene vector is shown here. 7-week-old wildtype C57BL6 mice were administered systemically via tail vein with AAV9s containing NTG or m31 gRNA at the indicated dose. 11 weeks post administration animals were sacrificed, tissues were collected for further analysis. (D) Levels of Lama1 mRNA was assessed by RT-qPCR normalized by Hprt. Relative Lama1 level is presented as fold change relative to AAV9-GNDM-NTG treated animals. Bars represent mean values and SEM is represented by error bars (n = 4). **P < 0.01, ****P < 0.0001, n.s. =not significant. Unpaired t-test. (E) A comparison of RNAseq reads alignments across the Lama1 gene between AAV9-GNDM-NTG and AAV9-GNDM-m31 treated gastrocnemius (GC) muscle. (F) Volcano plot comparing whole transcriptome RNA-seq data between AAV9-GNDM-m31 and AAV9-GNDM-NTG treated GC muscles. Red dots highlight Lama1 as the most significantly induced transcripts globally. The data are representative of the average of four independent biological replicates.

Optimization of the AAV vector for efficient epigenetic editing is crucial due to inherent packaging size limitation of the AAV genome. Following extensive optimization efforts of individual vector elements, we have developed a final vector with streamlined components to maximize efficiency and specificity. This construct included a CK8-based promoter for muscle-specific expression, a codon-optimized GNDM transgene for enhanced protein expression, a compact polyadenylation signal, and a gRNA m31 expression cassette, collectively called GNDM-m31 (Figure 1C).

7-week-old wild-type mice were injected with AAV9 encapsulating GNDM-m31 (AAV9-GNDM-m31) at a dose of 1.5 x 10^14^ vector genomes(vg) per kilogram (vg/kg). At 11 weeks post-administration, the animals were sacrificed, and various tissues were collected for qPCR analysis. Our results demonstrate that animals treated with AAV9-GNDM-m31 exhibited significant upregulation of Lama1 mRNA specifically in skeletal and cardiac muscle tissue, with no observed upregulation in non-target tissues such as the liver (Figure 1D).

To assess the specificity of AAV9-GNDM-m31 *in vivo*, total RNA from gastrocnemius muscle were isolated and whole transcriptome RNA sequencing analysis was performed. Mapping reads from AAV9-GNDM transduced tissues to the Lama1 gene revealed selective upregulation of Lama1 expression with m31 gRNA, which had no impact on the expression of genes located immediately upstream or downstream of the Lama1 gene. (Figure 1E). Global analysis of mRNA expression from NTG or m31 sgRNA transduced animals also show minimal impact on total gene expression, with Lama1 mRNA having uniquely high and significant expression (Figure 1F). These data highlight the transcriptional specificity of CRISPR-GNDM-m31. Along with its targeted delivery and expression in specific tissues, this underscores the potential of CRISPR-GNDM technology for precise gene therapy applications.

### AAV9-GNDM-m31 induces Lama1 protein expression in muscle tissues and alleviates pathophysiology in DyW disease model

Homozygous DyW mice are completely deficient in Lama2 protein, and exhibit severe disease phenotypes of LAMA2-CMD with premature death^28^. To assess the efficacy of AAV9-GNDM-m31 in the DyW disease model, postnatal day 2 (PND2) mice were treated with AAV9-GNDM-m31 or AAV9-GNDM-NTG at 4.5 x 10^11^ vg per animal (roughly 3 x 10^14^ vg/kg) via the temporal vein (Figure 2A). Consistent with previous observations with wildtype mice, Lama1 mRNA level is significantly induced in various skeletal muscle tissues in AAV9-GNDM-m31 treated DyW mice 4 weeks after injection (Figure 2B). In addition, substantial Lama1 protein expression was detected by immunofluorescent staining (Figure 2C). This expression pattern closely resembled that of Lama2 protein in wild-type mice, with Lama1 protein localizing at the basement membranes of muscle fibers and cardiomyocytes. These results suggest that the induced LAMA1 protein forms functional laminin complexes, and integrates effectively into the extracellular matrix of these tissues. DyW mice exhibit elevated serum creatine kinase (CK) levels, an indicator of muscle damage. When treated with AAV9-GNDM-m31, their serum CK levels significantly decreased (Figure 2D). Muscle morphology was also improved with AAV9-GNDM-m31 treatment. In AAV9-GNDM-NTG treated DyW mice, H&E staining revealed large groups of small-sized muscle fibers with many centrally localized nuclei, indicative of active muscle degeneration (Figure 2E). In contrast, muscles from mice treated with AAV9-GNDM-m31 exhibited near-normal morphology (Figure 2F) with significantly fewer central nuclei (Figure 2G), suggesting GNDM-induced Lama1 expression protected myofibers from degeneration.

**Figure 2:**
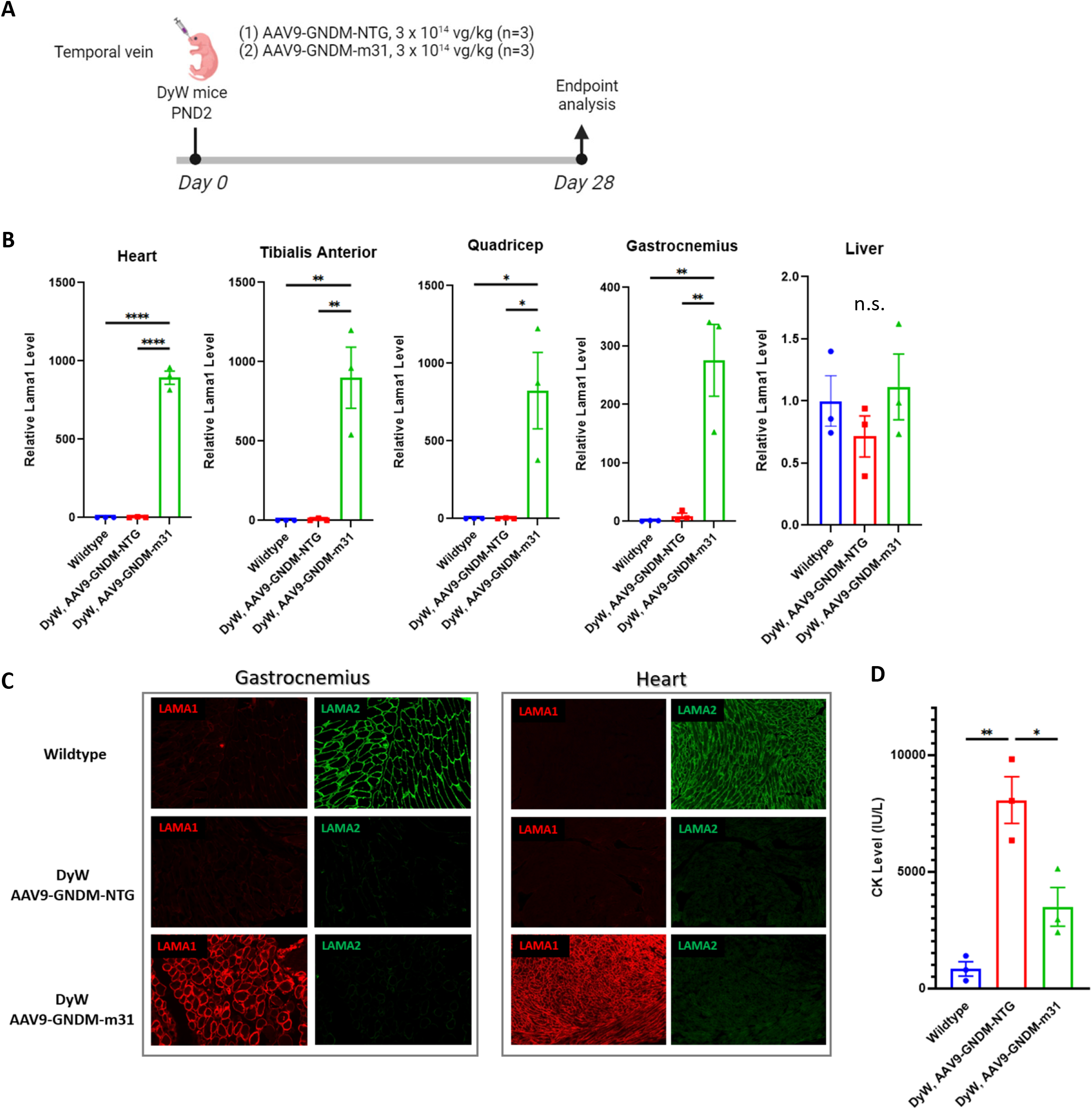
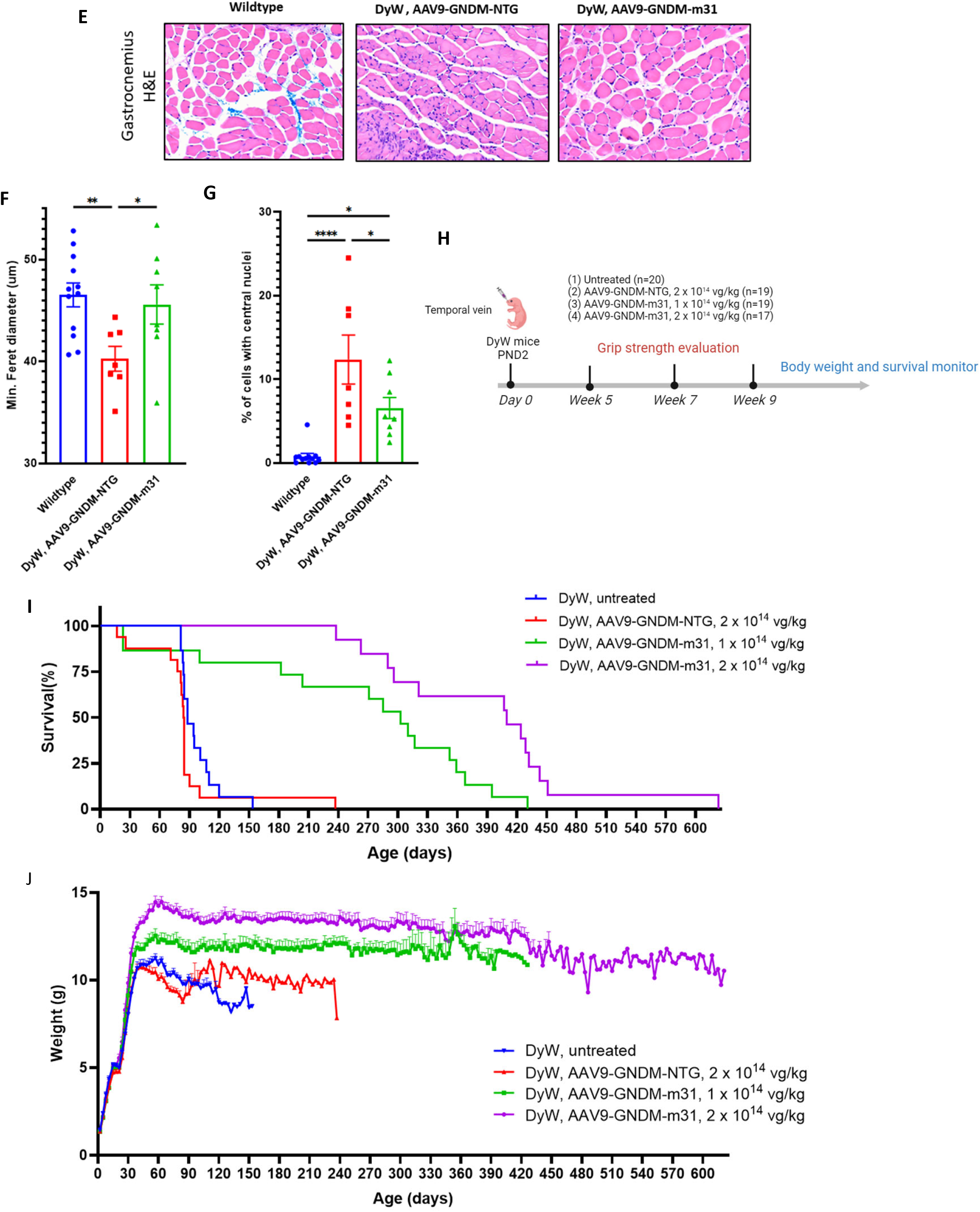
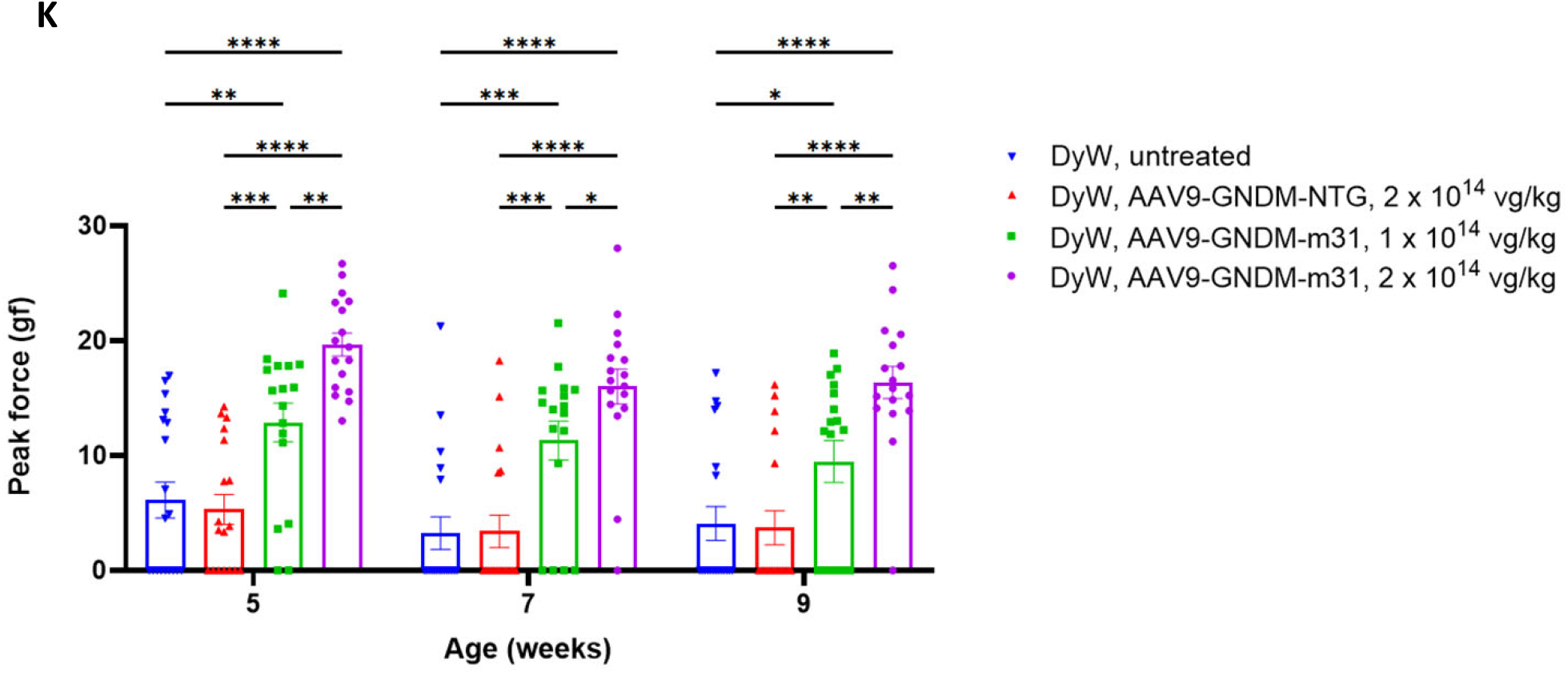
AAV9-GNDM-m31 upregulates Lama1 and significantly improves lifespan and ameliorates muscle pathology in DyW mice. (A) PND2 mice were administered systemically via temporal vein with AAV9-GNDM-NTG or m31 gRNA at the indicated dose. After 28 days, mice were euthanized and various tissues were harvested for endpoint analysis. (B) Levels of Lama1 mRNA was assessed by RT-qPCR and normalized to Hprt. Relative level of Lama1 was presented as fold change relative to age-matched wildtype C57BL6 mice. Bars represent mean values and SEM is represented by error bars (n = 3). *P < 0.05, **P < 0.01, ****P < 0.0001, n.s. =not significant. One-way ANOVA with Tukey test. (C) Immunofluorescence staining images of LAMA1 and LAMA2 in cross-sections of GC muscle (left) and heart (right) from each treatment group. (D) Relative CK levels in 4-week-old mice. Bars represent mean values and SEM is represented by error bars (n = 3). *P < 0.05, **P < 0.01. One-way ANOVA with Tukey test. (E) Representative haematoxylin and eosin (H&E) staining of GC muscles from each treatment group. (F) Muscle fiber size determined as minimum Feret’s diameter in GC muscle from each treatment group. (G) Percentage of myofibers with centrally located nuclei. (F-G) Bars represent mean values and SEM is represented by error bars. *P < 0.05, **P < 0.01, ***P < 0.001, ****P < 0.0001. One-way ANOVA with Tukey test. (H) PND2 mice were administered systemically via temporal vein with AAV9s containing NTG or m31 gRNA at the indicated dose. Animal survival was monitored daily and body weight was measured every two or three days. Grip strength was measured 5, 7 and 9 weeks post administration. Mice that died before wean were excluded from analysis. (I) Kaplan-Meier survival analysis of DyW mice of all treatment groups. ***P <0.001, log-rank test (AAV9-GNDM-m31, 1 x 10^14^ vg/kg or 2 x 10^14^ vg/kg versus untreated DyW or AAV9-GNDM-NTG, 2 x 10^14^ vg/kg). (J) Bi-weekly weights throughout the life span of DyW mice of all treatment groups. (K) Quantification of forelimb grip strength at 5, 7 and 9 weeks of age for each experiment group. Bars represent mean values and SEM is represented by error bars. *, p<0.05, **, p<0.01, ***, p<0.001, ****, p<0.0001. Two-Way ANOVA with Fisher’s LSD test.

We next investigated whether the observed histopathological improvements in DyW mice also led to an extended lifespan and enhanced growth. A large cohort of DyW mice were divided into four groups and received the following treatments at PND2: 1. Untreated; 2. AAV9-GNDM-NTG (2 x 10^14^ vg/kg); 3. AAV9-GNDM-m31 low dose (1 x 10^14^ vg/kg); and 4. AAV9-GNDM-m31 high dose (2 x 10^14^ vg/kg) (Figure 2H). After 80 days post-injection, 100% of the DyW mice treated with high doses of AAV9-GNDM-m31 were alive, compared to 50% survival in the untreated or NTG-treated groups. The survival improvement with treatment became more pronounced at around 120 days post-injection. At this point, DyW mice treated with AAV9-GNDM-m31 showed extended survival (80% in the low dose group and 100% in the high dose group), compared to only a 5% survival rate for untreated or AAV9-GNDM-NTG treated mice. Notably, over 50% of mice in the low dose group survived past 300 days, and over 400 days in the high dose group (Figure 2I). Additionally, mice treated with AAV9-GNDM-m31 gained significant body weight compared to untreated or AAV9-GNDM-NTG treated mice, which was maintained throughout their life, indicating durable therapeutic effect of AAV9-GNDM-m31(Figure 2J). We also measured muscle function at 5, 7, and 9 weeks of age. The DyW mice treated with AAV9-GNDM-m31 showed a significant improvement in grip strength at all experimental timepoints, in a dose-dependent manner (Figure 2K). Collectively, these results underscore the robust efficacy of GNDM technology for the muscle-related symptoms of LAMA2-CMD.

### Durable expression of GNDM and Lama1 up to 12 months with transient immune response against the transgene in Dy2J disease model

In order to evaluate the immune response and durability of the transgene expression in proinflammatory disease microenvironment, we used Dy2J mice. DyW model was not suitable for this purpose because of their short survival time precluding comparison against untreated cohort. Dy2J disease model has a near-normal life span, albeit milder phenotype. Systemic administration of AAV9-GNDM-m31 at 3 x 10^14^vg/kg in 6-week old Dy2J mice induced robust expression of GNDM and Lama1 in muscle tissue starting from week 1 to up to 1 year post-injection, the longest period tested (Figure 3A). Moreover, we noted sustained transgene expression lasting up to 16 months in wild-type mice injected with AAV9-GNDM-m31 (data not shown). Analysis of serum samples for antibody responses by IgG ELISA showed that AAV9-GNDM-m31 administration induced antibody responses to Cas9 (Figure 3B). Quantification of Cas9-specific T cell responses using interferon-gamma (IFNγ) ELISPOT assay showed a transient induction of T cell responses at week 2 which declined sharply by week 4 and remained very low throughout the study period (Figure 3C). These Cas9-specific T cell responses led to a partial decline in GNDM expression in muscle tissue by week 4. However, remaining GNDM expression persisted throughout the study period. Most importantly, the induced LAMA1 protein level remained unchanged for up to 12 months (Figure 3D). Further, we evaluated the effect of Cas9-specific immune responses on targeted muscle tissue. Immunohistochemistry analysis of gastrocnemius (GC) muscle tissue using CD8-specific and CD11b-specific antibodies showed increased CD8 T cell infiltration at week 2 after AAV9-GNDM-m31 administration which declined significantly following the week 4 time point and remained at low levels throughout the study period. On the other hand, the CD11b positive population was unchanged (Supplemental Figure 1). QPCR analyses of different immune markers in the muscle tissue revealed elevated CD8, IFNγ and Granzyme B levels at week 2 and followed a similar declining trend of T cell responses in ELISPOT data (Figure 3E). On the other hand, Foxp3 expression increased in AAV9-GNDM-m31 administered animals at week 2 and remained high until after 1 year. In agreement with previous reports, muscle targeted AAV administration induced persistent Foxp3 T cells in the muscle tissue^29^, indicating a counterbalance of immune response in the muscle tissue. These results indicate that, although AAV9-GNDM administration induces transient T cell and antibody responses to the transgene, they have minimal effect on transgene (GNDM) expression and no effect on target gene (Lama1) expression.

**Figure 3:**
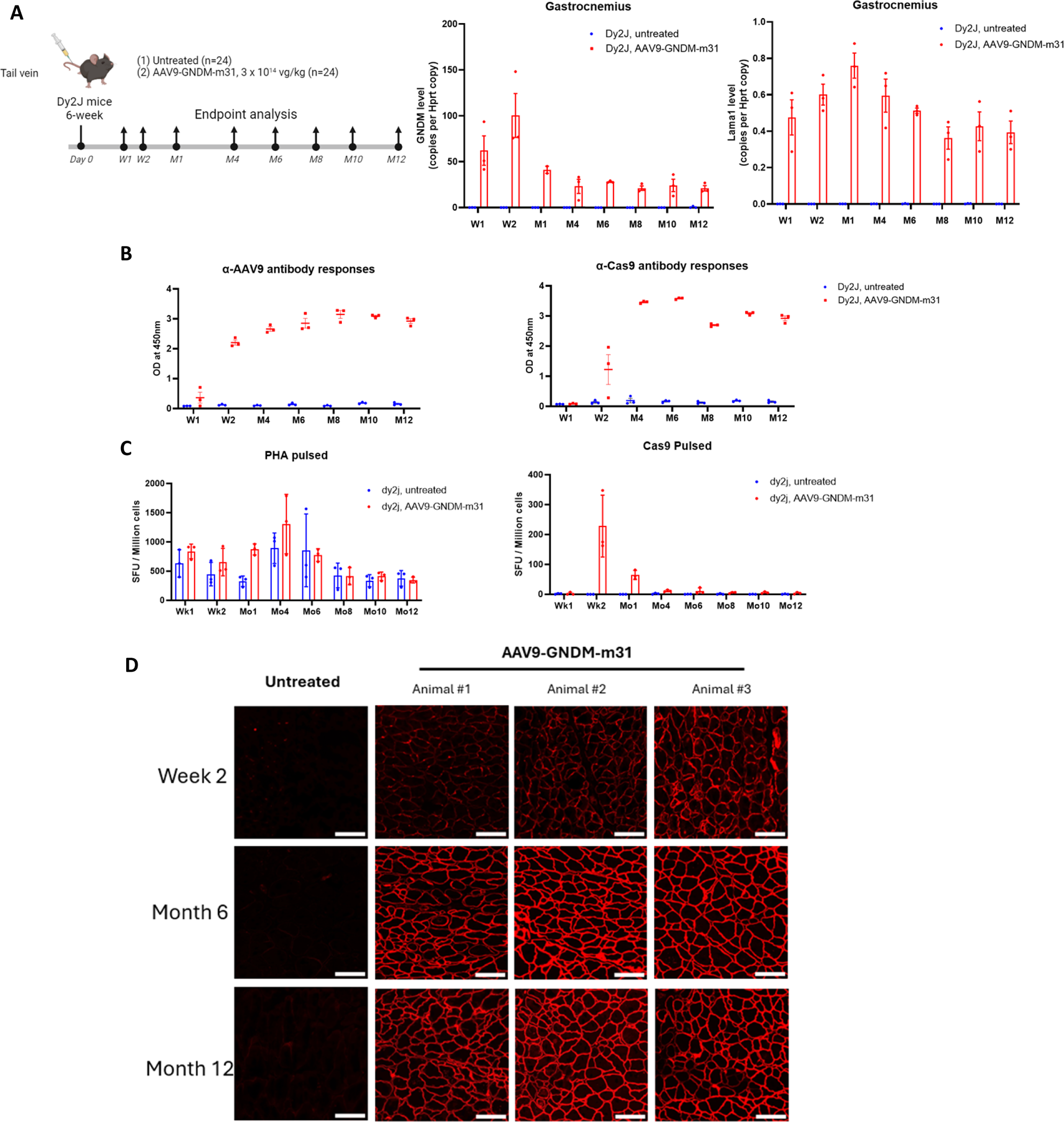
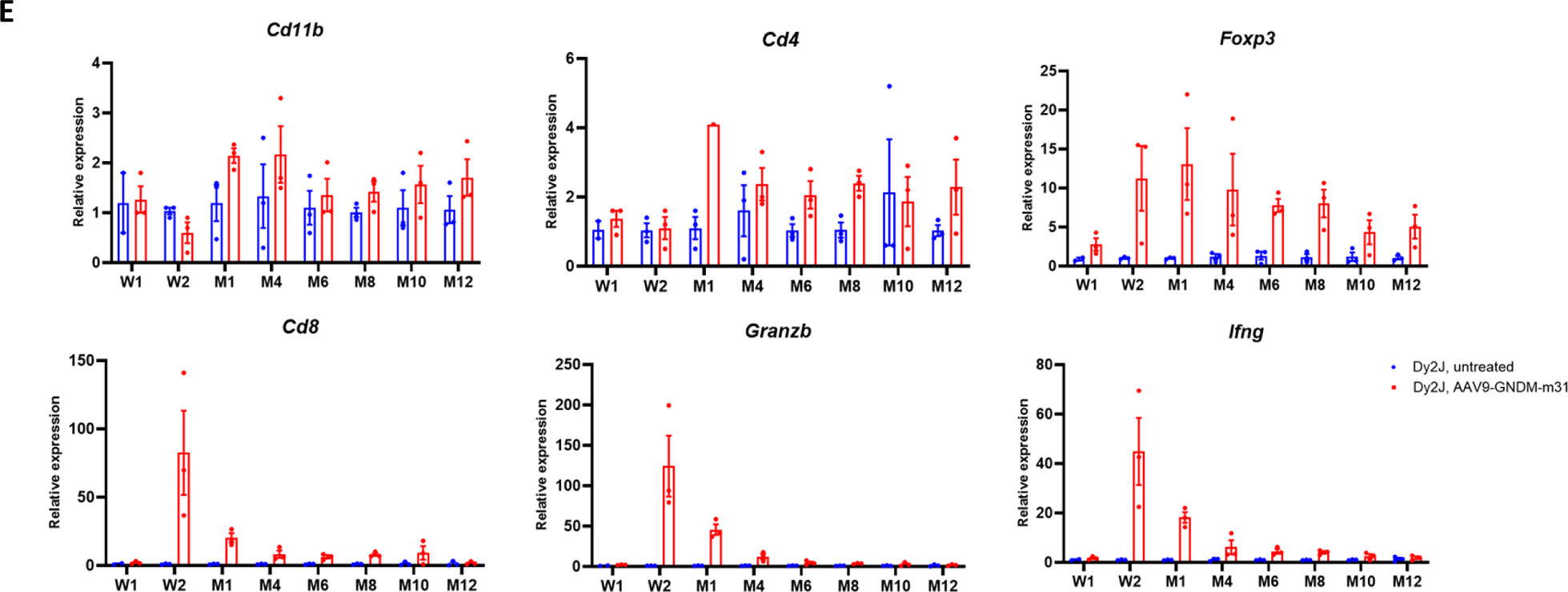
AAV9-GNDM-m31 induces durable expression of GNDM and LAMA1 in Dy2J mice up to 12 months but with only transient immune response against transgene. (A) Dy2J mice were administered AAV9-GNDM-m31 or left untreated for the study duration and sacrificed for analysis at indicated time point (n=3 per time point per treatment group). RNA was extracted from gastrocnemius tissue, and Lama1 and GNDM mRNA level is quantified by probe-based RT-ddPCR and presented as Hprt-normalized value. Bars represent mean values and SEM is represented by error bars. (B) Anti-Cas9 and Anti-AAV9 antibody responses in the serum were analyzed by ELISA and data are plotted as OD at 450 nm. (C) Cas9 specific T cell responses were analyzed at different time points by IFNg ELIPSOT assay. 200k splenocytes were re-stimulated with Cas9 protein (10ug/ml) or PHA (5ug/ml) for 48hrs and ELISPOT plate was developed. Data was presented as spot forming units for million splenocytes. No stimulation condition was used to subtract the background. (D) Immunofluorescence staining images of LAMA1 in cross-sections of the gastrocnemius from week 2, month 6 and month 12 timepoint (n=1 for untreated, n=3 for AAV9 treated). Scale bars, 100 µm. (E) RT-qPCR analysis of immune cell markers using cDNA derived from GC muscle. Gene expression was normalized by Hprt and presented as fold change relative to Dy2J, untreated samples. All values are obtained by averaging results from triplicates for each sample. Bars represent mean values and SEM is represented by error bars.

### Pronounced efficacy with MYOAAV at ten-times lower dose than AAV9 in DyW mice

In recent years, engineered myotropic AAV capsids have emerged as promising vectors for efficient gene payload delivery to muscle tissue^30^. MYOAAV is a modified version of AAV9 obtained through directed evolution, which involved adding a 7-mer peptide insertion containing the RGD sequence at the 588 position^31^. This modification significantly increases muscle tropism and potentially allows for the administration of viral vectors at much lower doses compared to conventional AAV9 vectors.

To evaluate the efficacy of MYOAAV-GNDM-m31, DyW mice were intravenously administered a dose of 2 x 10^13^ vg/kg at postnatal day 15 (PND15) (Figure 4A). This dose was significantly lower than that of AAV9-GNDM-m31. PND15 parallels a stage of human development of under 36 months, aligning better with our target LAMA2-CMD patient demographic for intervention. Mice were taken down at 6 weeks post-injection for analysis. Notably, the outcomes of this study were consistent with our previous findings utilizing AAV9 vectors. Specifically, we observed robust expression levels of the GNDM transgene and a significant upregulation of Lama1 mRNA exclusively within skeletal muscle, while non-targeting tissues such as the liver remained unaffected (Figures 4B, C, D). Immunofluorescent staining revealed significant LAMA1 protein expression at the basement membrane of muscle fibers (Supplemental Figure 2).

**Figure 4:**
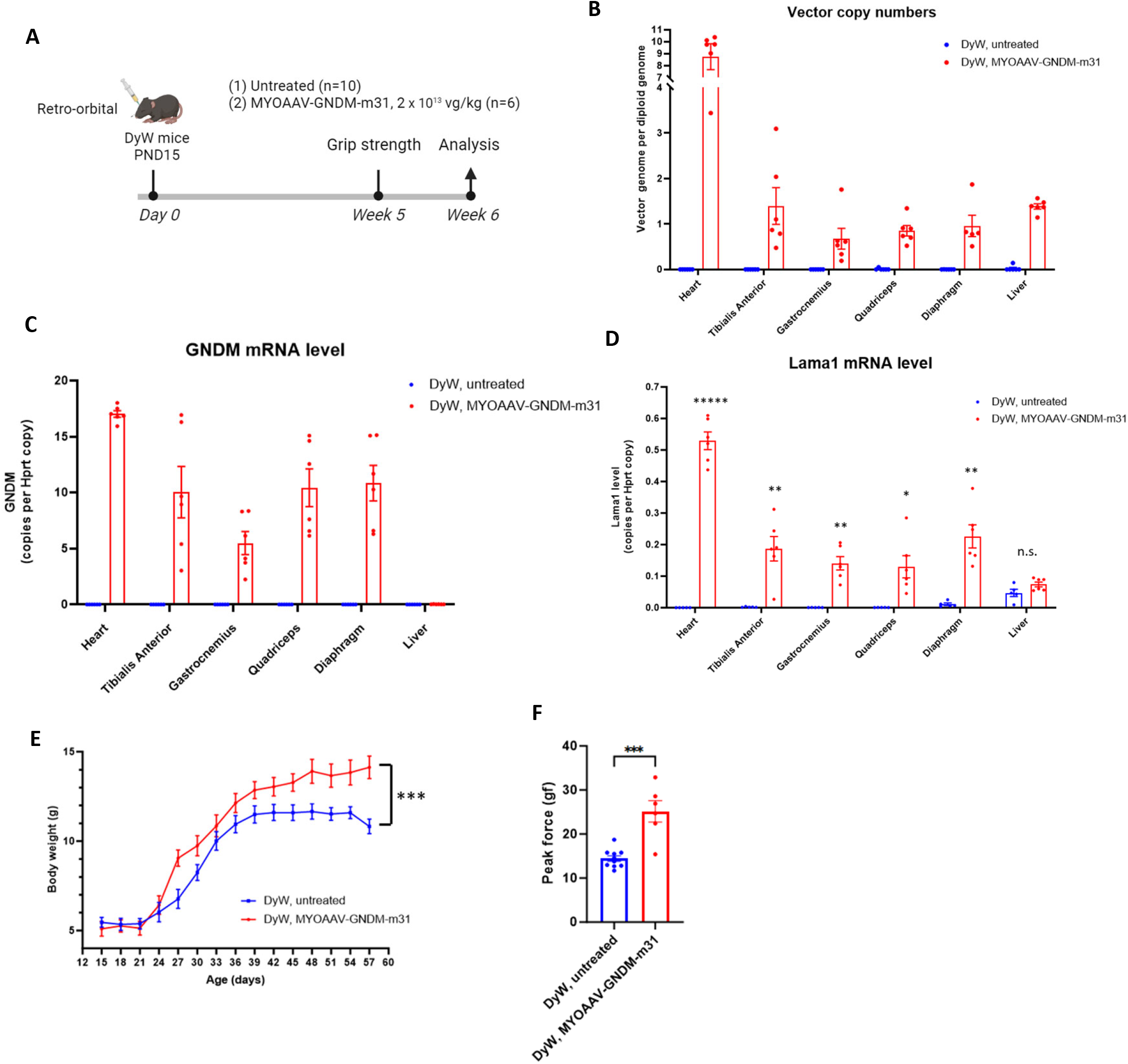
MYOAAV-GNDM-m31 demonstrated robust efficacy in DyW mice at significantly lower dose compared to AAV9. (A) DyW mice were administered MYOAAV-GNDM-m31 at PND15 or left untreated for the study duration. Body weights were measured bi-weekly. Grip strength was measured 5 weeks post-administration. One week later, mice were sacrificed, heart, liver, diaphragm, and skeletal muscle tissues were processed for mRNA and DNA analysis. (B) Vector copy number (VCN) was calculated by ddPCR copy number assay. (C)(D) Lama1 and GNDM mRNA levels were quantified by probe-based RT-ddPCR and presented as Hprt-normalized values. Bars represent mean values, and SEM is indicated by error bars. *, p<0.05, **, p<0.01, *****, p<0.00001, n.s. = not significant. Multiple unpaired t-test. (E) Body weight of 2- to 8-week old mice. Data are means ± SEM. ***P < 0.001, paired t-test. (F) Quantification of forelimb grip strength in 7-week-old mice. Peak force was measured as the maximal force applied before the animal released its forelimbs from the measurement apparatus while being pulled backwards by the experimenter. Data are means ± SEM. ***P < 0.0005, unpaired t-test.

Consistent with the previous findings, treated DyW mice demonstrated a notable increase in their body weights compared to untreated counterparts, indicative of enhanced overall health and muscle growth (Figure 4E). Furthermore, treated mice exhibited significantly improved grip strength, providing compelling evidence of enhanced muscle function following intervention (Figure 4F). Of particular significance is the fact that these results were achieved utilizing a dose that is more than five times lower than studies utilizing AAV9 vectors. This underscores the enhanced efficiency and potency of the novel MYOAAV capsid in targeting muscle tissue and its considerable advantage over conventional AAV9 vectors in gene therapy applications.

### Systemic administration of MYOAAV-GNDM-c58 in NHPs is safe and well-tolerated

Although epigenetic modulation has been demonstrated in mouse models with no apparent toxicity, there are higher safety and efficacy bars in taking the technology to patients. Particularly, concerns have been raised that systemically overexpressing foreign proteins such as Cas9 in primates could result in severe immune toxicity due to the complexity of their immune systems and outbread nature of the population. However, no studies have yet addressed these concerns in primates delivered AAV systemically with Cas9 epigenetic modifiers. Also, translation of AAV vector bioavailability from mice to large animals and humans remains to be established. Thus, we conducted a non-GLP study in adult cynomolgus macaques to evaluate the safety and biodistribution of our vector.

A similar *in vitro* gRNA screening was conducted to identify the most potent gRNA in primary cynomolgus myoblast cells, resulting in the selection of gRNA c58 for in vivo evaluation (Figure 5A). After one week of standard immunosuppressive treatment, three female cynomolgus macaques were administered MYOAAV-GNDM-c58 through peripheral vein injection at a dose of 1 x 10^14^ vg/kg (Figure 5B). (One animal was subsequently removed from further analysis due to retrospective identification of seroconversion to AAV positivity prior to dosing.)

**Figure 5:**
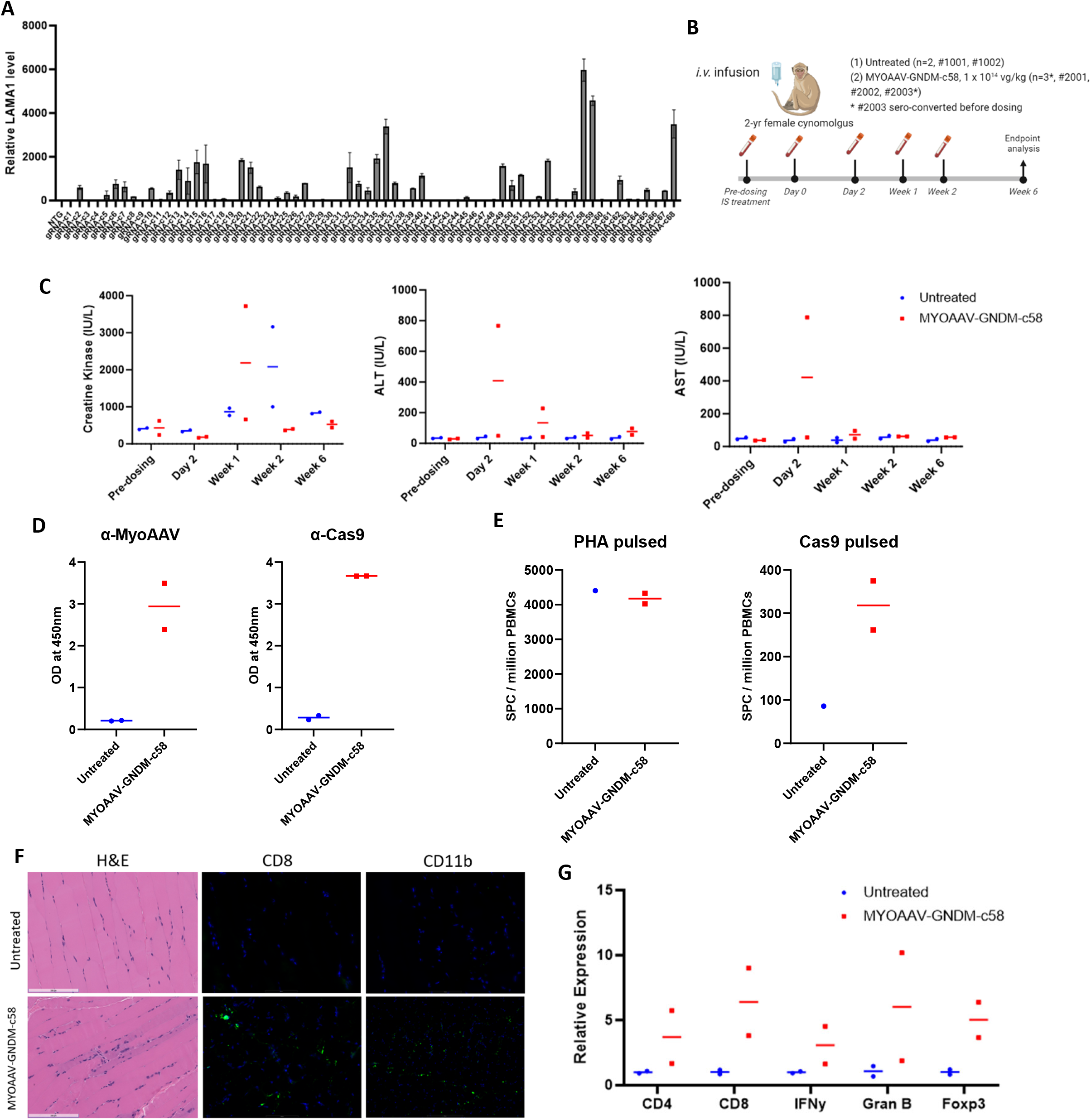
Systemic administration of MYOAAV-GNDM-c58 in non-human primates is safe and well-tolerated. (A) Summary of gRNA screening results using cynomolgus primary skeletal myoblast cells. Relative LAMA1 mRNA level is determined by qPCR where each sample expression values were normalized to the NTG sample values. Bars represent mean values and SEM is represented by error bars (n = 3). (B) MYOAAV-GNDM-c58 (1 x 10^14^ vg/kg) was administered by *i.v.* infusion to 2-year old cynomolgus macaques after standard immunosuppression(IS) treatment. Blood was sampled at indicated timepoint, and all animals were sacrificed 6 weeks after administration. Animal #2003 had AAV seroconversion before administration, thus excluded from analysis. (C) Quantification of serum CK, ALT and AST enzyme levels before and after systemic administration of MYOAAV-GNDM-c58. Data are presented as individual and mean value. (D) Anti-Cas9 and anti-MYOAAV antibody responses were analyzed by serum ELISA assay using plates coated with specific antigens and developed plates were scanned at 450nm. Data are presented as individual and mean OD at 450nm. (E) Cas9 specific T cell responses (IFNg ELISPOT assay) were analyzed at terminal time point. 125k PBMCs were re-stimulated with Cas9 protein (10ug/ml) or PHA (5ug/ml) for 48hrs and ELISPOT plate was developed. Data was presented as spot forming units per million PBMCs. No-stimulation condition was used to subtract the background. (F) GC tissues were sectioned for both H&E staining and immunofluorescent staining with anti-CD8 and anti-CD11b antibodies. Images were captured with ECHO Revolution color camera at 20x magnification. Scale bars, 100 µm scale. (G) RT-qPCR analysis of immune cell markers using cDNA derived from GC muscle. Gene expression was normalized by HPRT and presented as fold change relative to untreated animal samples. All values are obtained by averaging results from triplicates for each sample. Data are presented as individual and mean.

MYOAAV-GNDM-c58 was well tolerated with no obvious adverse events and no detectable changes in body weight and wellbeing of the animals throughout the study. Blood chemistry revealed only a temporary elevation of liver enzymes, which normalized within 2 to 3 weeks (Figure 5C). Other clinical pathology findings were comparable between treated and control animals, indicating a favorable safety profile of the vector. Serum antibodies against MYOAAV capsid and GNDM protein (Cas9) (Figure 5D), and T cell responses against Cas9 protein in the ELISPOT assay (Figure 5E) were observed 6 weeks after the injection; showing both humoral and cellular immune systems recognized GNDM protein and mounted responses. However, there was minimal lymphocyte infiltration in the target muscle tissues and no obvious tissue damage was observed (Figure 5F). QPCR analyses of different immune cell type markers in the muscle tissue revealed elevated CD4, CD8, IFNγ and Granzyme B levels, reflecting observed lymphocyte infiltration. However, Consistent with Dy2J data, Foxp3 expression also increased in MYOAAV-GNDM-c58 administered animals (Figure 5G), indicating a counter balance of immune response in the muscle tissue. These results support the safety of our MYOAAV-GNDM construct from an immunological perspective.

### Administration of MYOAAV-GNDM-c58 in adult NHPs showed favorable biodistribution with robust target engagement and safety

The ddPCR analysis of the AAV genome in a range of muscle and non-muscle tissues showed that MYOAAV-GNDM-c58 had widespread biodistribution throughout the body following a single administration (Figure 6A). The GNDM expression analysis revealed that a muscle-specific promoter used in the vector allowed for strong GNDM expression only in muscle but not in non-muscle tissues such as the liver (Figure 6B). Notably, MYOAAV-GNDM-c58 vector showed up to 60-fold enhanced GNDM mRNA expression in muscle tissues as compared to our historical datasets generated in NHPs with AAV9 vector (data not shown). These data indicate that despite the high expression of the transgene and the presence of immune responses against the protein, GNDM expression remained robustly high until the end of the study, 43 days post injection.

**Figure 6:**
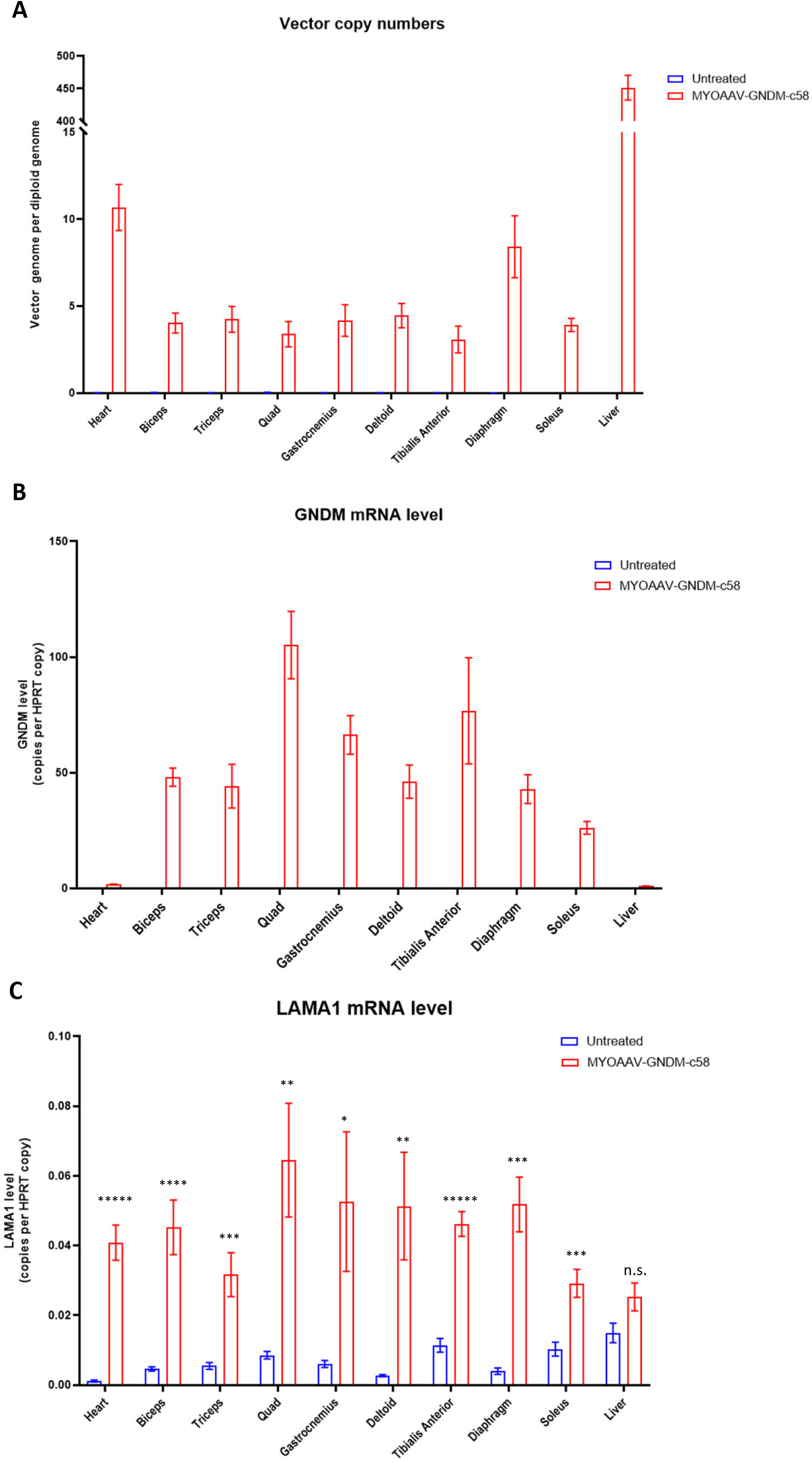
Administration of MYOAAV-GNDM-c58 in adult NHPs showed favorable biodistribution with robust target engagement. Multiple pieces of heart, liver, diaphragm, and skeletal muscle tissues per animal were processed for mRNA and DNA analysis. (A) Vector copy number (VCN) was calculated utilizing ddPCR copy number assay. (B) and (C) GNDM and LAMA1 mRNA levels were quantified by probe-based RT-ddPCR and presented as HPRT-normalized values. Bars represent mean values of tissue replicates from animals (n=2) per group, and SEM is indicated by error bars. *, p<0.05, **, p<0.01, ***, p<0.001, ****, p<0.0001, *****, p<0.00001, n.s. = not significant. Multiple unpaired t-test.

To demonstrate the induced LAMA1 mRNA levels in relation to a housekeeping gene, LAMA1 mRNA levels were normalized to endogenous HPRT mRNA levels. The induced LAMA1 levels were between ∼3-6 % relative to the endogenous HPRT levels in all muscle tissues analyzed (Figure 6C). As untreated animals otherwise have nearly undetectable levels of LAMA1 expression in muscle tissues, and given the high expression of HPRT gene, this represents a significant increase of LAMA1 mRNA. LAMA1/HPRT mRNA levels were also elevated in the heart (∼4%). The liver displayed some LAMA1 mRNA expression (∼3%), though not significantly above the endogenous level in the liver, indicating muscle-specific induction of LAMA1 with MYOAAV-GNDM-c58. This is a clear demonstration that the CRISPR-GNDM technology is able to reactivate gene expression from a physiologically silenced locus in large animals.

### Administration of MYOAAV-GNDM-c58 in juvenile NHPs is safe and well-tolerated

LAMA2-CMD is a congenital disorder and its symptoms manifest at birth. Patients initially follow a somewhat normal physiological development path and experience body and muscle growth. However, muscle wasting begins shortly after birth and early intervention is expected to yield superior clinical benefit^18^. Thus the target population of our treatment is anticipated to be infants and young children. We extended our studies to assess the safety and efficacy of MYOAAV-GNDM in 8-month-old juvenile NHPs (Figure 6A). This age group in NHPs closely resembles the developmental stage of our target patient demographic. We administered 5 x 10^13^ vg/kg of MYOAAV-GNDM-c58 to these juvenile NHPs, a dose carefully chosen to presumably strike an optimal balance between efficacy and safety for this younger and more vulnerable patient group. Also, we extended the study period to 13 weeks (as opposed to 6 weeks in adult NHP study), which allowed for a more thorough observation of the treatment’s effects over time, enhancing our understanding of the safety, efficacy and durability of the MYOAAV-GNDM treatment.

**Figure 6:**
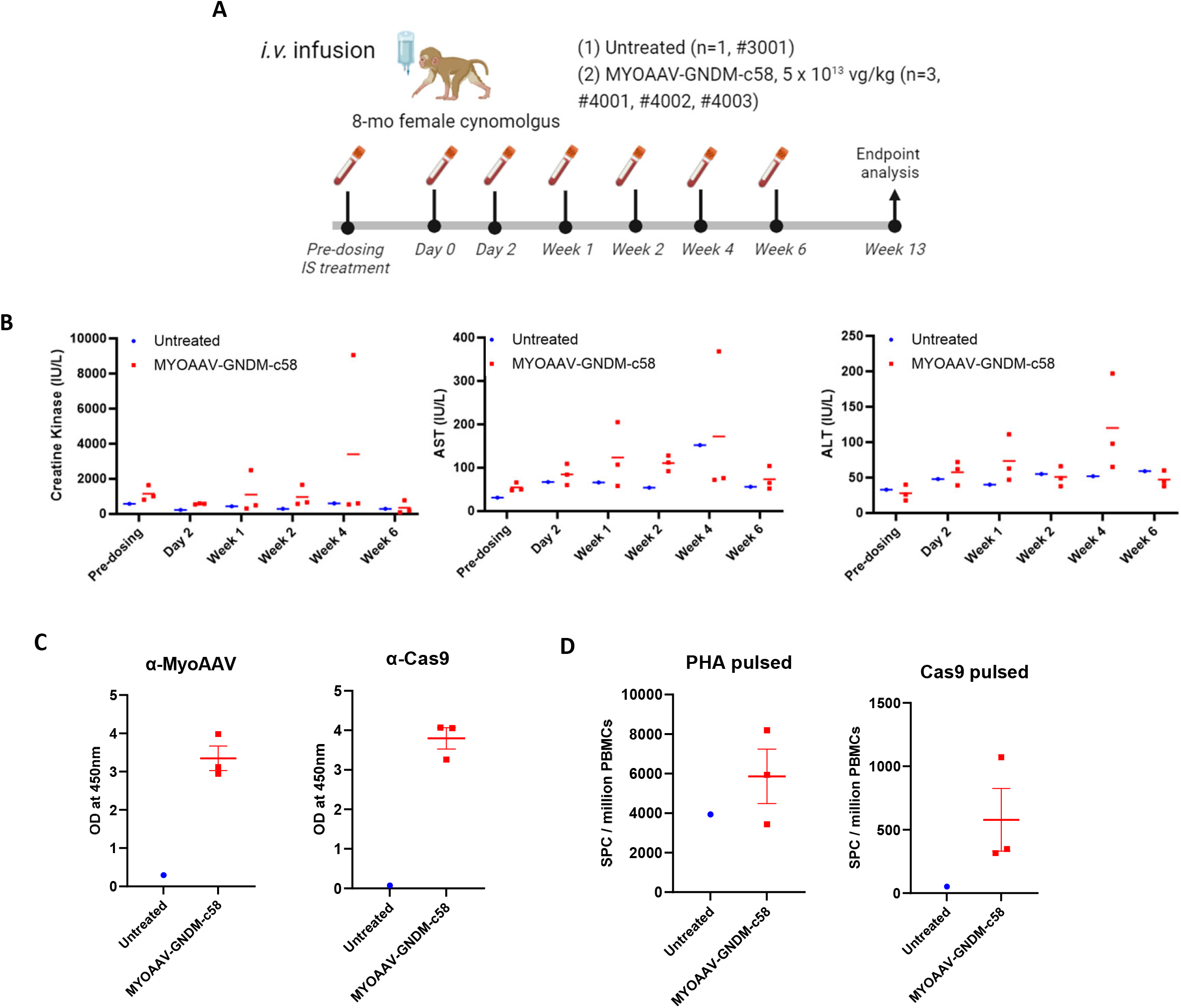
Administration of MYOAAV-GNDM-c58 in juvenile NHPs is safe and well-tolerated. (A) MYOAAV-GNDM-c58 (5 x 10^13^ vg/kg) was administered by *i.v.* infusion to 8-month old cynomolgus macaques after standard immunosuppression(IS) treatment. Blood was sampled at indicated timepoint, and all animals were sacrificed 13 weeks after administration. (B) Quantification of serum CK, ALT and AST enzyme levels before and after systemic administration of MYOAAV-GNDM-c58. Data are presented as individual and mean value. (C) Anti-Cas9 and anti-MYOAAV antibody responses were analyzed by serum ELISA assay using plates coated with specific antigens and developed plates were scanned at 450nm. Data are presented as individual and mean OD at 450nm.(D) Cas9 specific T cell responses (IFNg ELISPOT assay) were analyzed at terminal time point. 125k PBMCs were re-stimulated with Cas9 protein (10ug/ml) or PHA (5ug/ml) for 48hrs and ELISPOT plate was developed. Data was presented as spot forming units per million PBMCs. No-stimulation condition was used to subtract the background.

Similar to our prior study with 2-year-old NHPs, all juvenile animals in this extended 13-week study tolerated the MYOAAV-GNDM treatment well, with no observed toxicity (Figure 6B). Histopathological evaluation revealed no adverse macroscopic findings in the tissues examined. Microscopic examination showed minimal mononuclear cell infiltration in the biceps brachii and gastrocnemius muscle, which was likely due to mechanical injury rather than the treatment. All other microscopic findings were considered unrelated to the administration of AAV, as they occurred at a low incidence/severity, were present in the control group, or were consistent with expected background findings in Cynomolgus macaques. These observations further corroborate the treatment’s safety profile.

### MYOAAV-GNDM-c58 in juvenile NHPs demonstrate a superior pharmacodynamic profile compared to adults

Vector copy numbers (VCN) analysis from various muscle tissues in adult and juvenile NHPs showed nearly half the number of VCN in juvenile NHPs (Figure 7A). This aligns with the halved dose used in the juvenile study. Surprisingly, even with this lower dose, the mRNA levels of the GNDM transgene in juvenile NHPs were significantly higher compared to those in older NHPs across all muscle tissue types (Figure 7B). This indicates a more efficient GNDM transgene expression in juvenile NHPs. Additionally, higher expression of the GNDM transgene leads to significantly increased induction of LAMA1 mRNA (Figure 7C). It should be noted that the duration of the two NHP studies were different which may also play a role in the superior pharmacodynamic profile in the juvenile monkeys. However, these findings suggest the potential for increased efficacy with greater safety at lower dosages in younger subjects.

**Figure 7:**
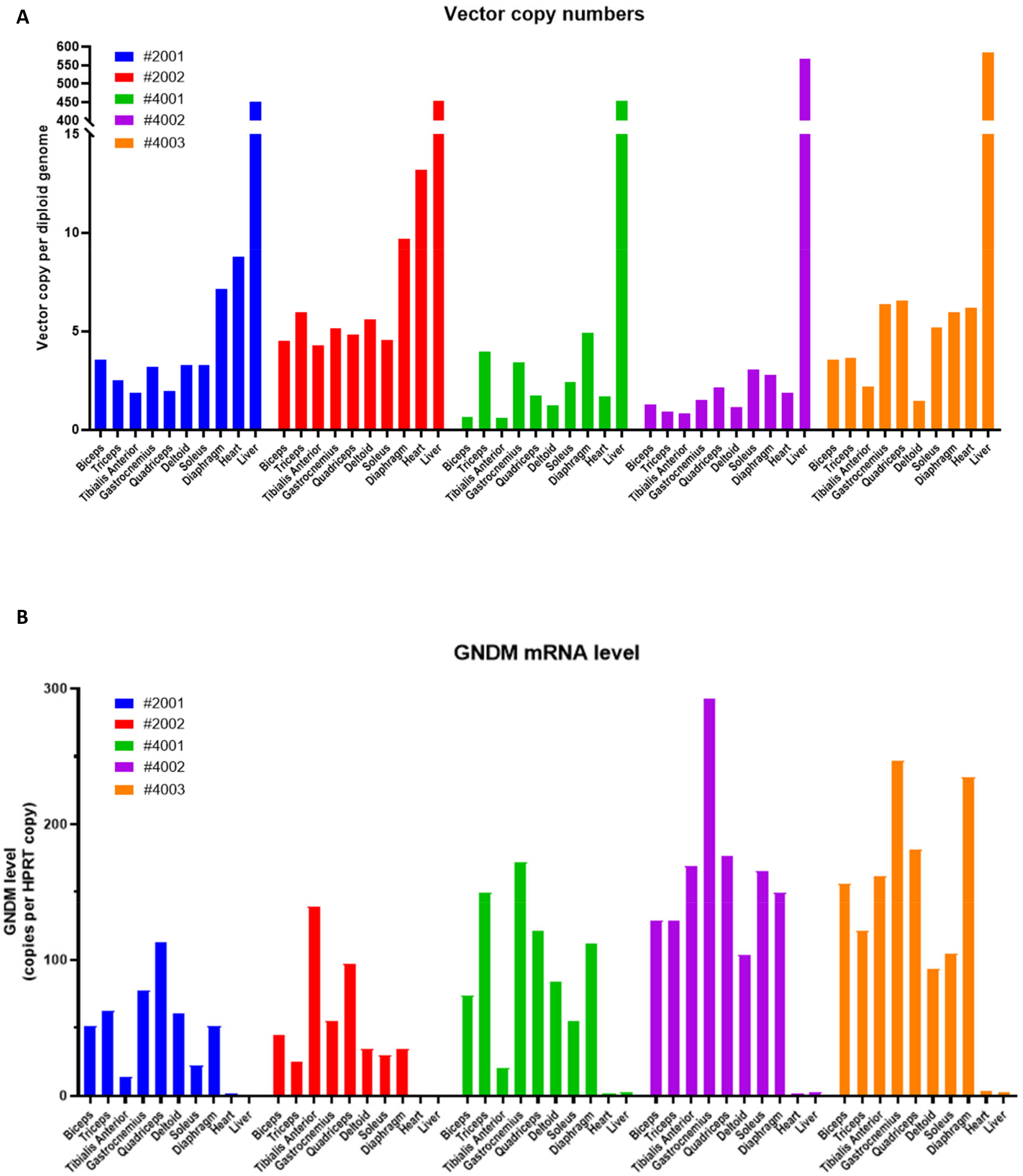
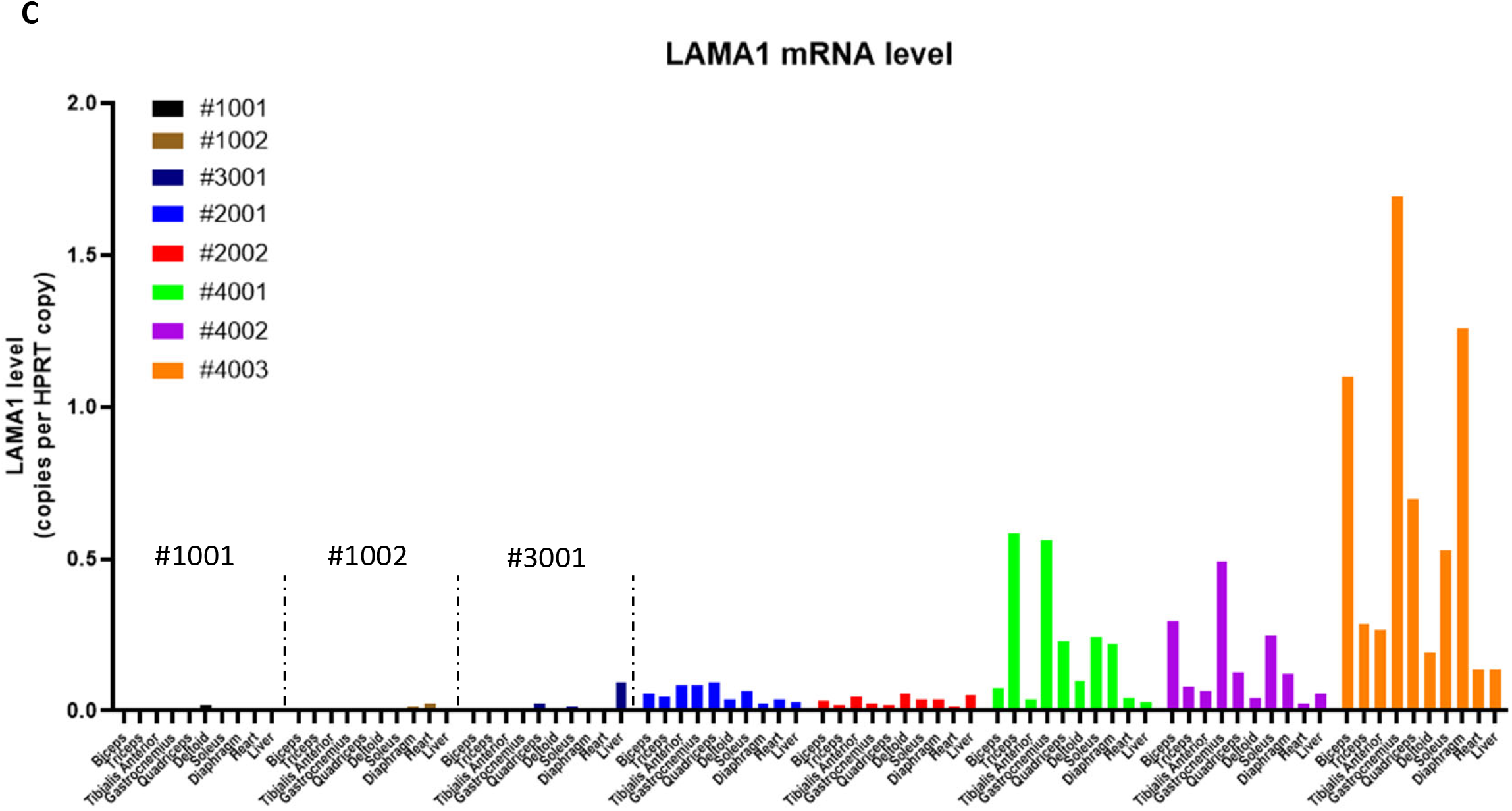
The lower dose of MYOAAV-GNDM-c58 in juvenile NHPs demonstrate a superior pharmacodynamic profile. Multiple pieces of heart, liver, diaphragm, and skeletal muscle tissues per animal were processed for mRNA and DNA analysis. (A) Vector copy number (VCN) was calculated utilizing ddPCR copy number assay. (B) and (C) GNDM and LAMA1 mRNA levels were quantified by probe-based RT-ddPCR and presented as HPRT-normalized values. Bars represent mean values of tissue replicates from individual animal.

## Discussion

While there have been reports in small animals using experimental systems, it has never been addressed whether epigenetic activation approaches such as ours in a viable therapeutic format could induce a robust target gene expression from silenced loci of large animal’s genome. Our CRISPR-GNDM^®^ technology has taken the CRISPRa system to a level more applicable for a viable therapeutic by improving several key points. First, the entire CRISPRa component was packaged into a single AAV vector, eliminating the need for two AAV vectors. Second, the single gRNA was chosen for the maximal potency of LAMA1 upregulation and minimum off-target gene expression changes. Third, a potent activation moiety for LAMA1 upregulation was fused to the dCas9. Fourth, the tissue-specific promoter was used to drive GNDM expression specifically in muscle tissues, avoiding off-target tissue expression of GNDM. Finally, the use of a muscle tropic capsid significantly enhanced AAV transduction efficiency into muscle tissues, lowering overall dose and vector-related toxicity. Using this vector, we have demonstrated the success of this approach in rodent and NHP models, marking a pioneering achievement in the use of a single AAV vector-based therapy for a condition akin to LAMA2-CMD.

LAMA2-CMD is a rare, congenital autosomal recessive muscular dystrophy, affecting 8.3 in 1 million individuals and accounts for one third of all CMDs. The most common cause of death in early-onset LAMA2-CMD is respiratory tract infection, with 30% of patients dying within the first decade of life^18^. To date, there is no effective treatment available for LAMA2-CMD patients and current treatment options focus only on symptom relief.

Although the most straightforward strategy to treat loss-of-function type genetic disorders is to replace the defective gene, the size of LAMA2 coding sequence precludes an AAV-mediated supplement approach. Several groups have attempted to address this challenge. One approach is to use smaller linker proteins, such as aLNNd and mini-agrin^32, 33^, to restore connections typically established by Laminin 211, by effectively utilizing Laminin-411. However, this approach employs two AAV vectors and is not practical for the current gene therapy product development. Despite not being expressed in muscle tissues, laminin-α1 is the most structurally and functionally similar protein to laminin-α2 and is known to compensate for the loss of laminin-α2 in the LAMA2-CMD disease setting. Thus, multiple groups have explored leveraging laminin-α1 to compensate for the loss of laminin-α2. An administration of Laminin-111 protein complex was reported to be effective in ameliorating disease phenotypes in the DyW mouse model^23^. However, a more durable approach is to use epigenome editing technology to upregulate the endogenous LAMA1 gene. *Kemaladewi et al.* have successfully demonstrated amelioration of the disease phenotype in the Dy2J mouse model^26^, however, using a two-AAV vector system with three gRNAs. Our success in the disease models and preclinical testing with a single AAV vector not only validates our approach but also paves the way for application in human patients.

Our study is the first demonstration of MYOAAV for a therapeutic testing in NHPs. In addition to exhibiting the reported muscle tropism^31^, our study revealed favorable biodistribution of MYOAAV in juvenile NHPs. The mechanism for this finding is yet to be determined, however, the dose of 5 x 10^13^ vg/kg is extremely low compared to doses used in other systemic gene therapies for muscle disorders (e.g. 2∼3 x 10^14^ vg/kg), potentially highlighting the superior benefit of MYOAAV for muscle disease indications in pediatrics.

Gene therapy products often come with two concerns: capsid-related toxicity and transgene-mediated immunogenicity. Our MYOAAV-GNDM vector is made of novel muscle-tropic capsid with Cas9-containing transgene, and understanding the safety profiles of our novel vector is crucial for therapeutic application. Our study showed that transient elevation of liver enzymes and transient decline of platelets and RBC, indicating minimal/transient effect of MYOAAV capsid mediated innate immune responses. The kinetics and magnitude of the antibody response against MYOAAV capsid were similar to those reported against natural AAV capsids such as AAV9, relieving potential concerns raised against this novel capsid. Despite high hopes for CRISPR-Cas technology in addressing various genetic disorders with unmet medical needs, concerns have been raised against the usage of Cas9 protein for its exogenous and bacterially-derived aspects to be highly immunogenic. Previous studies conducted in mice and short-term study in Duchenne muscular dystrophy (DMD) dogs have shown conflicting results regarding the impact of Cas9-mediated immune responses on targeted tissue and overall safety. Some studies indicated no adverse effect^34^, while another demonstrated the induction of Cas9-mediated immune responses leading to the elimination of Cas9-expressing cells^35^. This disparity could be attributed to the specific animal model used in the studies and the prior exposure to *Staphylococcus aureus* from which Cas9 has derived. These animals may have presumably enhanced immune responses to Cas9^36^, as opposed to our NHPs pre-screened for Cas9-naive animals.

Our study was the first to address this concern in the relevant disease model and in NHPs. In mice, we utilized the Dy2J mice for its similarity in disease-genetics and proinflammatory tissue microenvironment to assess Cas9-mediated immunogenicity and its impact on pharmacodynamic and long-term therapeutic efficacy of our vector. Contrary to the study in a canine DMD model^34^, we observe the prolonged expression of GNDM and LAMA1 up to the maximum period tested (1 year), despite the mounted immune response against Cas9 protein. Our kinetic analysis of T cell subset gene expressions and tissue infiltrating immune cells show that the transient cytotoxic T cell infiltration was swiftly dampened followed by the continuous presence of Foxp3-expressing regulatory T cells otherwise not present in the pre-dosed muscle tissue. This observation reveals a hitherto unexpected role of regulatory T cells in supporting the sustained expression of gene therapy products expressing Cas9.

Furthermore, we evaluated the safety and immunogenicity of our AAV-GNDM therapeutic in NHPs. Consistent with the results from the Dy2J mouse study, there was only mild immune cell infiltration into the targeted tissues without any detectable histological alterations in dosed NHPs despite mounted humoral and cellular immune response. This is consistent with other reports, showing increased Foxp3 expression in targeted muscle tissue in both NHPs^37^ and humans^29^. In humans, Wagner et al. demonstrated that Foxp3 Tregs in human PBMCs, pre-exposed to *S. pyogenes*, effectively inhibit effector T cell function in-vitro^38^. Moreover, a recent study utilizing AAV9-Cas9 to edit latent SIV provirus in NHPs showed that a systemic administration was well tolerated without inducing any tissue damage^39^. These findings support that systemic delivery of Cas9 through AAV is a viable approach for therapeutic purposes.

The robust and persistent Cas9 expression in our study was somewhat surprising in the light of conventional immunology where the thesis is that exogenous antigens must be eliminated. Several speculations can be made: 1) an immune-tolerant nature of muscle tissue^37^, 2) continuous exposure to the transgene inducing T cell exhaustion^40^. Overall, our data demonstrates that AAV-GNDM administration established a balanced immune environment in the muscle tissue by Foxp3 T cell infiltration which modulates the immune mediated effect. The long-term impact of GNDM mediated immune responses in larger animals will be addressed in subsequent studies. The clinical strategy of our therapeutics would involve pre-screening of patients for pre-exposure to Cas9. Additionally, controlling initial immune response using potent combinations of immunosuppressive agents can induce tolerance to a transgene that is expressed at very high levels. Rituximab, prednisolone, and sirolimus have been successfully used in NHPs^41^ and AAV gene therapy clinical trials^42^. Similar combinations of immunosuppressive agents can be used prophylactically to control the initial wave of immune response and to induce transgene specific immune tolerance.

In conclusion, our study represents a significant step forward in demonstrating epigenetic activation for the treatment of LAMA2-CMD. By leveraging single vector GNDM technology, addressing the challenges of epigenetic gene activation, and introducing a novel capsid for enhanced delivery, we pave the way for an effective approach to treating certain debilitating conditions. Our study establishes a framework to employ epigenetic activation not only to treat LAMA2-CMD, but also for the treatment of various disorders where upregulating endogenous genes would bring clinical benefit.

## Method

### Cell culture

C2C12 cell line (ATCC) was maintained in DMEM (Invitrogen) containing 10% fetal bovine serum (FBS) and antibiotics (penicillin/streptomycin) at 37 °C with 5% CO2. For the transfection experiments, 12-well plates were seeded with approximately 100,000 cells per well to achieve 70%–80% confluency. Cells were transfected using 1 µg of plasmid DNA and 2 µL of Lipofectamine 2000 (Invitrogen) following the manufacturer’s instructions. Successful transfection was assessed by cell survival following 48 hours of selection with 2 µg/mL puromycin, after which RNA was harvested for gene expression analysis.

### Lentivirus generation and transduction

HEK293TA cells were seeded at 750,000 cells per well in 6 well cell culture dishes in 2 ml growth medium (DMEM media supplemented with 10% FBS and 2 mM fresh L-glutamine, 1 mM sodium pyruvate and non-essential amino acids) and incubated at 37°C/5% CO_2_ for 24 hours. The next day TransIT-VirusGEN transfection reactions were set up according to manufacturer’s protocol with 1.5 µg packaging plasmid mix (1µg pCMV delta R8.2 and 0.5 µg pCMV-VSV-G) and 1 µg of transfer plasmid containing sequence encoding GNDM and indicated sgRNAs. Lentivirus was harvested 48 hours following transfection by passing media supernatant through a 0.45 µM PES filter (VWR). Until ready to use, the purified and aliquoted lentiviruses were stored in −80°C freezer.

Primary cynomolgus macaque skeletal muscle myoblast cells were obtained from BioIVT. Cells were cultured in SkGM-2 Skeletal Muscle Growth BulletKit medium (Lonza). For transduction, cells were seeded at 125,000 cells/well in 12 well plates containing growth medium and incubated at 37°C/5% CO_2_ for 24 hours. The next day, lentivirus supernatant supplemented with 8 µg/ml Polybrene (Sigma) was added to each well. To enhance transduction efficiency, cells were centrifuged at 1200g for 1 hour before viral media was removed and replaced with fresh growth medium. 72 hours after transduction, cells were fed fresh 0.5 µg/ml puromycin selection medium every 2-3 days. Following 7-10 days of cells being in selection medium, cells were harvested and RNA extracted with RNeasy 96 kit (Qiagen) as directed by the manufacturer.

### RNA and DNA extraction from animal tissues

To isolate mRNA from mouse tissue, a hybrid protocol using TRIzol™-chloroform phase separation and the Qiagen RNeasy 96 (Qiagen) was used. First, the tissue was lysed in lysing matrix A tubes (MP Biomedicals) with 1 mL of TRIzol™ (Invitrogen) in a bead mill. After centrifugation for 10 minutes at 12,000 x g, the supernatant was transferred to Phasemaker™ tubes (Invitrogen) and 160 µL of chloroform was added. Tubes were vigorously hand shaken for 20 seconds and then centrifuged at 4C for 18 minutes at 10,000 x g for phase separation. Next, the aqueous supernatant was mixed with an equal volume of 70% ethanol. The mixture was then transferred to a Qiagen RNeasy 96 plate and the isolation was continued according to the manufacturer’s protocol.

Genomic DNA isolation was carried out by following the Qiagen DNeasy Blood and Tissue Kit (Qiagen). Briefly, tissue was lysed overnight in buffer ATL with Proteinase K and moderate agitation at 56°C. The lysates were mixed with buffer AL/E and transferred to a DNeasy 96 plate where, after centrifugation, columns were washed with buffer AW1 and then AW2. Buffer AE was added to elute genomic DNA.

### Gene expression analysis by QPCR

cDNA was generated from no more than 1µg of total RNA according to High-Capacity cDNA Reverse Transcription Kit (ThermoFisher) protocol in a 10 µl volume. cDNA was then diluted 10-fold and analyzed using Taqman Fast Advanced Master Mix according to manufacturer’s instructions. Taqman probe-based real-time PCR reactions were processed and analyzed by QuantStudio 5 Real-Time PCR system.

For each sample, deltaCt values were calculated by subtracting the average Ct values of the target gene (derived from 3 technical replicates) from those of the housekeeping gene. Expression values for each sample were determined using the formula 2^-(deltaCt). These sample expression values were then normalized to the control sample expression values of each experiment to ascertain the relative expression of the target gene.

### RNA and VCN quantification by ddPCR

mRNA isolated from animal tissues were reverse transcribed using a High-Capacity cDNA Reverse Transcription Kit (ThermoFisher) following the manufacturer protocol. The resulting cDNA was used in Droplet Digital PCR (ddPCR) analysis to assess target gene expression using Hprt as the housekeeping reference. Isolated genomic DNA was used to assess vector copy number using Tfrc (mouse) or Tert (cynomolgus) as the reference, with a genomic copy number of two.

All ddPCR was carried out using the QX200 Automated Droplet Generator and QX200 Droplet Digital PCR System (Bio-Rad) for dual-color analysis according to the manufacturer’s instructions. Reaction mixtures consisting of the appropriate primer-probe sets, RNase-free water, ddPCR Supermix (Bio-Rad), and diluted cDNA or gDNA were made and loaded in the QX200 AutoDG to partition DNA into droplets. DNA was then amplified by PCR and droplets were individually counted and assessed for positive or negative FAM and VIC signals in the QX200 Droplet Reader.

Target gene expression was analyzed by dividing the number of positive FAM droplets (target gene) by the number of positive VIC droplets (housekeeper gene) for each sample. For genomic DNA analysis, the number of positive FAM droplets was divided by the number of positive VIC droplets and multiplied by 2 to determine the vector copy number per diploid genome.

### Mice and injections

Neonate PND2, juvenile PND15 and adult 7-12 weeks old mice were used for the study. C57BL/6J (Jax#000664) mice were used as wild type controls. Disease models DyW (Jax#013786) and Dy2J (Jax#000524) were cryo recovered by Jackson Labs and experimental mice were generated by inhouse breeding. Mice were given access to regular chow and water ad libitum. DyW mice were also given access to nutragel complete nutrition gels starting at wean. All studies were approved by the Institutional animal care and use committee. PND2 mice were injected via temporal vein (50 µL/mouse), PND15 mice were injected via retro orbital injection (10µL/g), and adult mice were injected via tail vein injection (10µL/g).

### NHP and injections

Twelve healthy female two year old cynomolgus monkeys were enrolled in the first arm of the study. A second arm included four eight and nine-month old juvenile (N=3 male and N=1 female) cynomolgus monkeys. Animals were confirmed seronegative for relevant AAV serotype and Cas9 prior to enrollment. All enrolled animals were treated with immunosuppression (Methylprednisolone, 20mg, IM) one day prior to AAV dosing and once weekly throughout the duration of the in-life period. All treatment animals were anesthetized and AAV was infused intravenously over 30 minutes. Animals were closely monitored post injection for any acute reaction. At the close of the in-life period, animals were anesthetized and terminal blood was collected. Animals were humanely euthanized and necropsy and tissue collection of various organs was performed.

All animal care and protocols were performed and approved under relevant Institutional Animal Care and Use Committee guidelines.

### AAV production and purification

Suspension HEK293 cells were transfected with plasmids necessary for AAV production – an Ad helper plasmid, GNDM gene of interest plasmid, and a rep/cap plasmid. Three days post-transfection cells were lysed, treated with Benzonase (MilliporeSigma), and clarified through depth filtration and sterile filtration. Clarified lysates were purified over affinity chromatography and Anion-Exchange (AEX) chromatography for enrichment of genome containing capsids before final concentration and buffer exchange.

### Immunofluorescence and H&E staining

OCT-embedded mouse muscle tissue was sectioned at 7 µm on glass slides and a hydrophobic barrier was drawn around the tissue with a PAP pen (Vector Labs). The tissue was then rehydrated in PBS and subsequently fixed at 4⁰C for 8 minutes with 4% Formal Fixx (Epredia). Slides were then washed in PBS and permeabilized at room temperature for 10 minutes in a solution of 0.2% Triton in TBS (50 mM Tris pH 7.4, 90 mM NaCl). Next, the sections were blocked for 1 hour at room temperature with 10% normal goat serum (Invitrogen). Slides were incubated overnight at 4⁰C with primary antibody in 10% normal goat serum with 1% bovine serum albumin (Fisher Scientific, BP1600-100). The next morning, slides were washed in TBS and incubated for 1 hour at room temperature with secondary antibodies. Slides were washed in TBS and then mounted and cover slipped with an antifade mounting medium with DAPI (Vector Labs).

Primary antibodies used in this study include: rabbit monoclonal anti-mouse laminin-α1 (Custom made by Genscript, 1:200); rat monoclonal anti-mouse laminin-α2 (4H8-2, Santa Cruz Biotechnology, 1:50); rat monoclonal anti-CD8 (clone 4SM16, Invitrogen, 1:200); anti-CD11b (M1/70 Cat# 14-0112-82, eBiosciences). All Photomicrographs were taken with ECHO Revolution color camera (ECHO revolution microscope, USA), using the ECHO Application Suite software (Version 2).

### SaCas9 enzyme linked immune absorbent spot (ELISpot) assay

Frequencies of SaCas9 specific T cells were determined using the IFN-γ ELISpot assay. Mouse splenocytes were isolated from the spleens of control and treated mice at each time point and 200k splenocytes seeded into each well. Cells were stimulated in vitro for 48 h with 10 mg/ml of SaCas9 protein (Genscript Inc.). IFN-γ producing cells were indirectly detected as spots using the mouse-specific IFN-γ ELISpot kit (Immunospot, Cleveland OH, USA) as per the manufacturer’s instructions. Spots were analyzed and counted using the ImmunoSpot software (Version 7, Cellular Technology Limited, Cleaveland, OH, USA) on the ImmunoSpot® S6 Analyzer (Cellular Technology Limited). The number of cells producing IFN-γ was presented as a spot forming unit (SFU) per million cells. PHA was used as the positive control in each assay. Media was used as the negative control in each assay. Untreated naïve mice were evaluated as the baseline. Assays were performed in triplicates.

### Monkey IFNγ ELISPOT assay

PBMCs were isolated from each treated and control animal and 25k PBMCs were seeded into each well and stimulated in vitro for 48 h with 10 mg/ml of SaCas9 protein (Genscript Inc.). IFN-γ producing cells were indirectly detected as spots using the Monkey-specific IFN-γ ELISpot kit (MabTech, USA) as per the manufacturer’s instructions. Spots were analyzed and counted using the ImmunoSpot software (Version 7, Cellular Technology Limited, Cleveland, OH, USA) on the ImmunoSpot® S6 Analyzer (Cellular Technology Limited). The number of cells producing IFN-γ was presented as spot forming unit (SFU) per million cells. PHA was used as the positive control in each assay. Media was used as the negative control in each assay. Untreated naïve mice were evaluated as the baseline. Assays were performed in duplicates.

### Antibody assay

Serum anti-AAV9 or anti-SaCas9 antibody levels were evaluated using an enzyme-linked immunosorbent assay (ELISA). Briefly, 2.5 ng/ml of the AAV9 or SaCas9 protein (Genscript Inc, USA) in carbonate buffer was coated on a microtiter plate. The serum was underwent five-fold serial dilution starting at 1:25 dilution. Horseradish peroxidase (HRP) conjugated goat anti-mouse IgG was used (1:10000; Southern Biotech, USA) for mice serum samples and goat anti-Cyno IgG for monkey serum samples (Southern Biotech, USA) to detect anti-Cas9 IgG. The absorbance of each sample was measured at 450 nm in the GloMax Microplate Spectrophotometer (Promega, USA). Each sample was run in triplicate and data was plotted as absorbance value at 450 nm vs 1:100 serum dilution.

### RNA sequencing

Illumina sequencing was performed by GeneWiz, LLC, where gene expression libraries were prepared using the NEBNext Ultra RNA Library Prep Kit (NEB, Ipswich, MA, USA) according to the manufacturer’s protocol. Sequencing libraries were clustered on three lanes of an Illumina HiSeq flow cell and sequenced using a 2×150 Paired End configuration. Resulting raw sequence data (.bcl files) was converted to fastq files and demultiplexed using Illumina’s bcl2fastq 2.17 software, where one mismatch was allowed for index sequence identification. Fastq files were aligned to human genome assembly GRCh38.p12, which contains the modification of GNDM sequence, using the STAR aligner^43^. Differential analysis was conducted using DESeq2^44^ and plots were generated with plotly (https://plot.ly) using custom R scripts. Additionally read were mapped and visualized on the genome using Integrated Genomics Viewer (https://igv.org/)

### Statistical analyses

Statistical analyses for every experiment were performed using GraphPad Prism software version 10. Analyses methods were described in the figure legends.

---

**Supplemental Figure 1:**
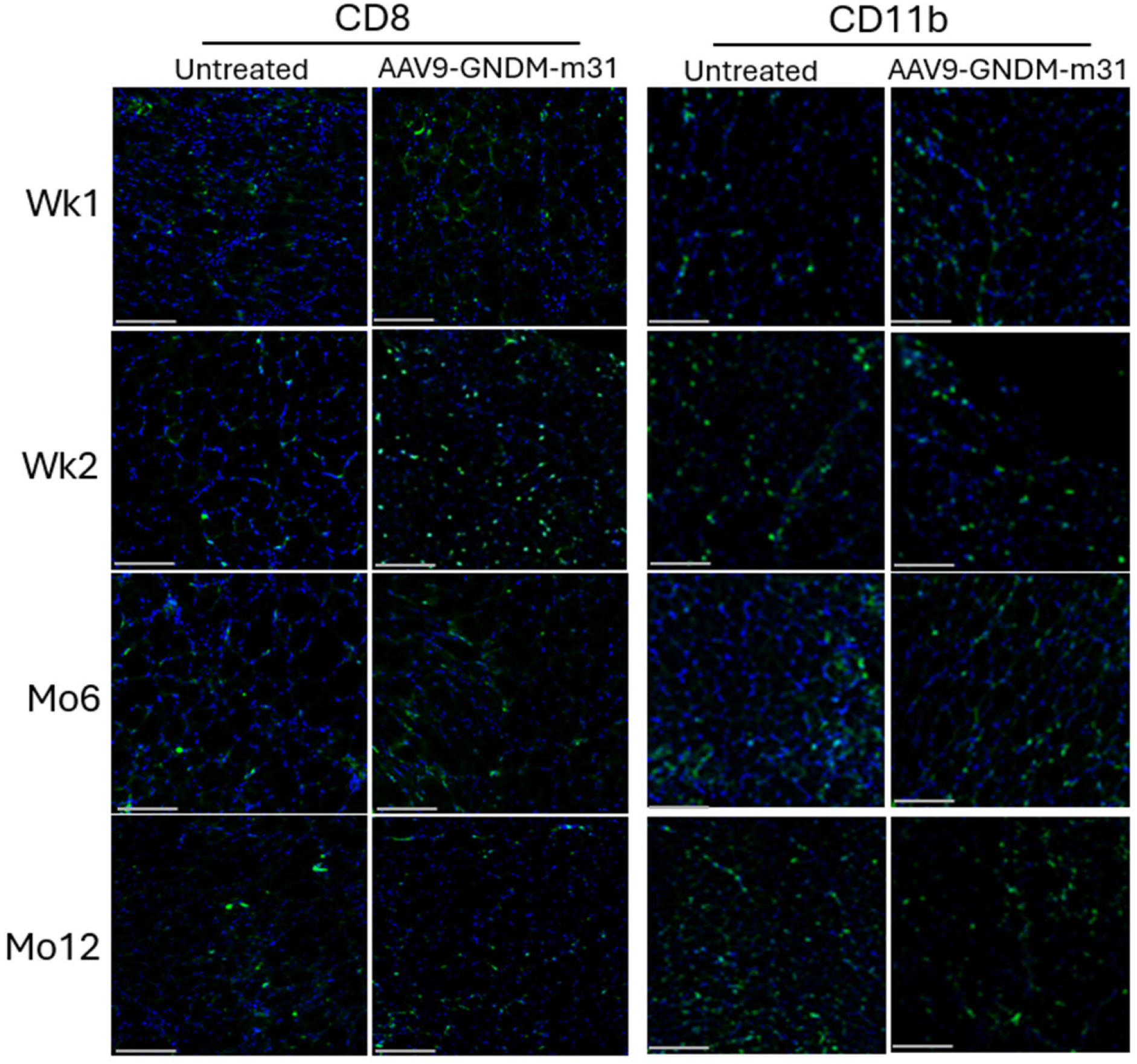
AAV9-GNDM-m31 systemic administration induces transient elevation of infiltrated immune cells in the muscle tissue. GC muscle from Dy2J mice at indicated time points was processed for OCT sections. OCT sections were stained with anti-CD8 and anti-CD11b rat monoclonal antibodies and probed with anti-rat AF-488 secondary antibody. Scale bars, 100 µm.

**Supplemental Figure 2:**
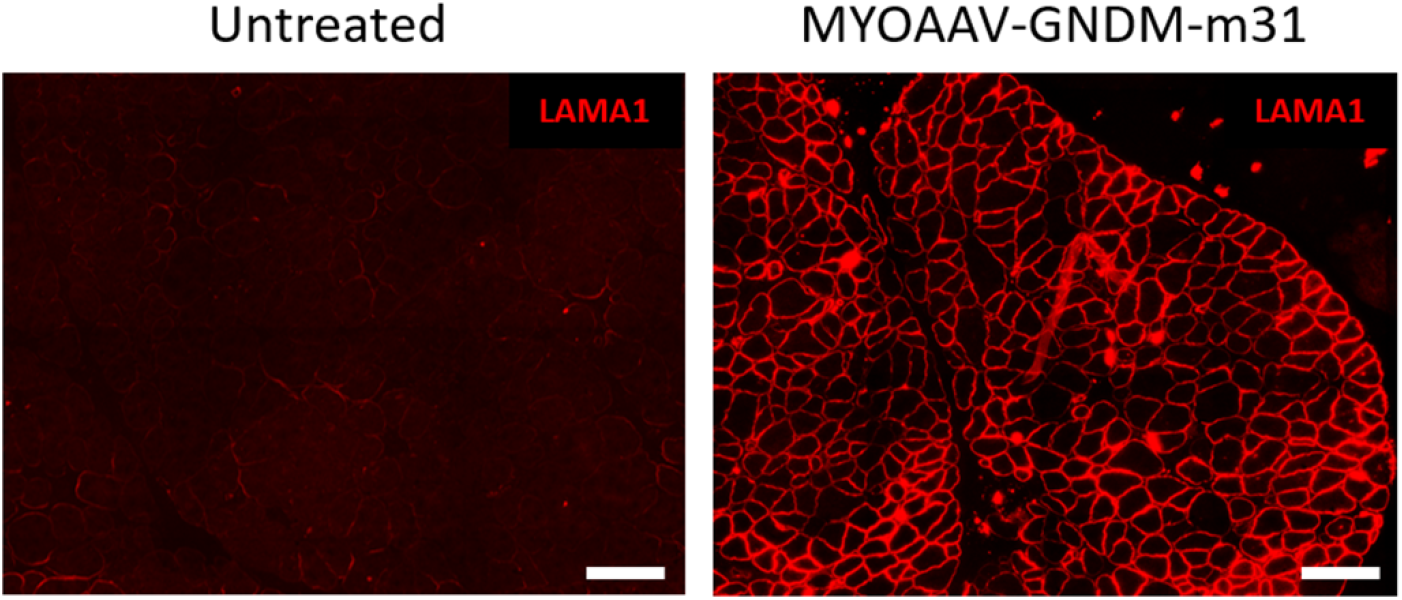
Expression and localization of induced LAMA1 protein in MYOAAV-GNDM-m31 treated DyW GC muscle. Immunofluorescence staining of LAMA1 in cross-sections of DyW GC muscle. Scale bars, 100 µm.

